# A pilot study of stable isotope fractionation in *Bombyx mori* rearing

**DOI:** 10.1101/2023.01.08.523185

**Authors:** Li Hao, Yujie He, Jinzhong Lu, Liling Jia, Yong Liu, Dan Yang, Shuai Shao, Gang Lv, Hailiang Yang, Hailing Zheng, Xuhong Cui, Yang Zhou, Zhiqin Peng

## Abstract

Hydrogen, oxygen, carbon, and nitrogen isotopes derived from three different strains of silkworms at different life stages involved in silkworm rearing, were measured to understand the fractionation characteristics of stable isotopes at different stages of silkworm development, and to trace the movement of these isotopes from food to larva to excrement and finally to silk. We found that silkworm strain had little effect on δ^2^H, δ^18^O and δ^13^C values. However, a large difference was found in the δ^15^N levels of newly-hatched silkworms between Jingsong Haoyue and Hua Kang No.3 orthogonal strains, suggesting that the mating and egg laying differences may result in an inconsistent kinetic nitrogen isotope fractionation. The δ^13^C values of silkworm pupae and silkworm cocoon also displayed significant differences, suggesting that heavy carbon isotopes are greatly fractionated from the larva to the silk during cocoon formation. Overall, these results may be used to clarify the relationship between isotope fractionation and the ecological process of the *Bombyx mori* and expand our ability to resolve stable isotope anomalies at a small regional-scale level.

## Introduction

The *Bombyx mori* is an important silk producer, which is believed to have been domesticated from the wild mulberry silkworm 5000-10000 years ago (Goldsmith et al., 2005). The composition and quality of the silk produced is dependent on the raw material of the cocoon, which in turn greatly depends on the conditions during the silkworm rearing stages. Silkworms undergo complete metamorphosis during rearing. The four developmental stages of egg, larva, pupae, and adult are completely different in terms of morphology and physiology. After hatching, the larvae exclusively feed on mulberry leaves and develop into mature silkworms after approximately 20 days, prior to which they will pass through five instars. The first to third instar are referred to as the young larval stage, while the fourth and fifth instar compose the old larval stage (Dong et al., 2013). The ideal rearing conditions and the demand for mulberry leaves vary between these stages. This complex life history can be analyzed through stable isotope technology. Stable isotope technology is widely used in feeding ecology to study the fractionation mechanisms of stable isotopes, in different organisms or within different tissues of the same organism, that can correlate with various ecological factors (Hood-Nowotny and Knols, 2007; Hyodo, 2015; Morra et al., 2021). The fractionation and enrichment characteristics of ^13^C and ^15^N can be used to determine an animal’s food source (Chávez-Solís et al., 2020), study its nutritional health (Burian et al., 2020; Ishikawa et al., 2020) and explore complex life history processes (Ishikawa et al., 2020). Stable isotope values differ between consumers and their food sources by a relatively definite discriminant value.

While most researchers using stable isotope technology in feeding ecology have focused on aquatic systems (Belle et al., 2020; Gilbert et al., 2020; Reis et al., 2020), stable isotope technology has also been widely applied in geographical origin traceability of agricultural products. Regina et al (Knaller and Strbele, 2014) measured the stable isotope ratios for hydrogen, carbon, nitrogen, and oxygen of silk fabrics samples found in the underground palace of the Famen Temple, Shaanxi, China, to determine if stable isotope ratios can assist in provenance analysis of silk. Although data analyses were lacking in their report, the authors claimed that their results indicate that the silk raw material or silk cocoons of different provenance are clearly distinguishable from each other. They believed that through the combination of information regarding centers of sericulture during the Tang Dynasty, it may be possible to determine the region from which the silk fiber came.

Therefore, by exploring the isotope relationship in sericulture (production of silk fiber from the rearing of mulberry groves to the harvesting of cocoons), it is possible to trace the provenance of silk contained in both fabrics and cocoons. The first factor that must be accounted for when studying the origin of silk fabrics is that different breeds or rearing conditions may affect silkworm physiological development and cocooning. Secondly, the fractionation of stable isotopes is often associated with significant physiological shifts including changes in metabolic rate, energy expenditure, and fasting, depletion of fat reserves, and variation in hormones (Gannes et al., 1998). Previous studies provide compelling evidence that the magnitude of dietary fractionation differs among closely related species (Focken and Becker, 1998) and is influenced by various exogenous factors, such as starvation (Marshall D and McCue, 2008). Nearly all the research into the biochemical processes that govern incorporation of stable isotope ratio into consumer tissues stems from controlled feeding experiments in the lab(Forbes, 2006; Kling et al., 1992; Webb et al., 1998), while relatively little has been explored regarding the mulberry-silkworm-cocoon ecosystem. Importantly, given that the *Bombyx mori* exclusively feed on mulberry leaves, this provides the unique opportunity to investigate isotopic changes associated with metamorphosis and physiological processes, with little influence from diet. In theory, any isotopic offsets in mulberry-silkworm-cocoon ecological processes are the results of physiological effects associated with metamorphosis or any laboratory control conditions.

In this study, we used stable isotope analyses to determine the hydrogen, oxygen, carbon, and nitrogen isotope values of different silkworm tissues from different silkworm strains at different stages of development and measured the changes in stable isotope values of silkworms under starvation stress. Establishing a biochemical framework for the fractionation of stable isotopes using the mulberry-silkworm-cocoon ecosystem will expand our ability to resolve stable isotope anomalies in small regional-scale systems and potentially unlock new applications for stable isotope data in tracing the origin of silk textiles.

## Materials and Methods

### Silkworm rearing and sample preparation

Three *Bombyx mori* strains, Jingsong haoyue (JSHY), Haoyue Jingsong (HYJS), and Hua Kang No.3 orthogonal (HK3), which were bred by Zhejiang Academy of Agricultural Science, were selected for experiments. Larvae were reared on fresh mulberry leaves in an environment with a temperature of 25 ± 2 °C and relative humidity of 75% ± 4%. Fresh mulberry leaves, newly hatched silkworms, silkworms at the fifth instar, silkworm excrement of fifth-instar larvae, and cocoons were collected and dried at 70 □. To eliminate individual differences, all the samples were ground together into a fine powder.

### Starvation Experiment

Silkworms of JSHY and HYJS were chosen for a starvation experiment at the beginning of their third instar. For comparison, the larvae of each strain were divided into 7 groups and starved for 6, 12, 24, 36, 48, 60, and 72 hours, respectively. Following starvation, around 20 larvae were collected from each group, dried, and ground into powder for testing, while the remaining larvae were left to resume feeding.

#### Stable Isotope Analysis

Silkworm samples were transferred into a tin capsule for carbon and nitrogen isotope ratio analysis using a stable isotope ratio mass spectrometer (IRMS, MAT-235, Thermo Fisher Scientific Inc, USA) equipped with an elemental analyzer (Flash 2000 HT; Thermo Fisher Scientific Inc, USA). The samples were combusted at 960 □in a combustion reactor and a column oven at 50 □. A helium carrier gas with a purity of 99.999 % was set to 100 mL/min. When measuring the stable isotopes of hydrogen and oxygen, the tin cup was replaced with a silver cup and the samples were analyzed in a reactor tube at 1380 □.

The isotope ratios of H, O, C and N were expressed as parts per thousand (‰) against the international reference standards:

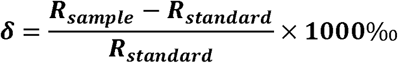

where R_sample_ represents the ratio of the heavy isotopes to the light isotopes of the element (C/N/H/O) in the sample, such as ^13^C/^12^C, ^15^N/^14^N, ^2^H/^1^H, ^18^O/^16^O; and R_standard_ is the ratio of the heavy isotopes to the light isotopes in the internationally recognized standard, using Vienna standard mean ocean water for H and O, Vienna Peedee Belemnite for C and N. The analytical precision and reproducibility (n=5; represented as mean ± standard deviation) based on the method validation analysis results were ≤ 0.1‰ for δ^13^C, ≤ 0.1‰ for δ^13^N, ≤ 0.4‰ for δ^2^H and ≤ 0.1‰ for δ18O, respectively. For the calibration, the certified reference materials IAEA-601 (δ^18^O=23.3‰), IAEA-CH-7 (δ^2^H=−100.3‰; δ^13^C =32.151‰) and IAEA-310 (δ^15^N = 47.2 ‰) were measured at intervals of 8 samples.

### Data Processing Method

Simple data distribution and mapping were performed with Excel (Microsoft) and Origin (OriginLab Corporation) 2018. Linear Discriminant Analysis (LDA) of stable isotope characteristic values in the different samples was performed using SPSS v22.0 (IBM).

## Results and discussion

### The effects of strains on the isotope composition ratios during silkworm development

The results of 3 different strains *Bombyx mori* isotope value are shown in Figure 1. The finding of a negligible difference in the value of δ^13^C among the newly-hatched silkworm and among fifth instar silkworms would indicate that silkworm strain has little effect on carbon stable isotope levels in silkworm rearing. In other words, the fractionation of δ^13^C by three strains *Bombyx mori* was consistent at the newly-hatched and fifth instar stages. However, our data showed a significant variation in δ^13^C values among silkworm pupae and cocoons, which might be driven by a differential selection between heavy and light elements during pupation and cocooning. The isotope analyses suggest that the molecular mechanisms underpinning the development of the silk glands lead to differences in nutrient allocation and ultimately results in variations in δ^13^C values between silkworm pupae and cocoons. Specifically, the process of silk protein production leads to the loss of protein, while lipids and carbohydrates are stored in silkworm pupae for metamorphosis(Sato and Azuma, 2016). Lipids and carbohydrates are known to have vastly different δ^13^C values compared to proteins (David et al., 2007; Hoering, 1981; Robbins et al., 2009; Terwilliger et al., 2002). Furthermore, there was also a difference in δ^13^C values among the 3 different strains silkworm pupae, which might be due to strain-specific differences in metabolic turnover rates.

**Figure 1.**
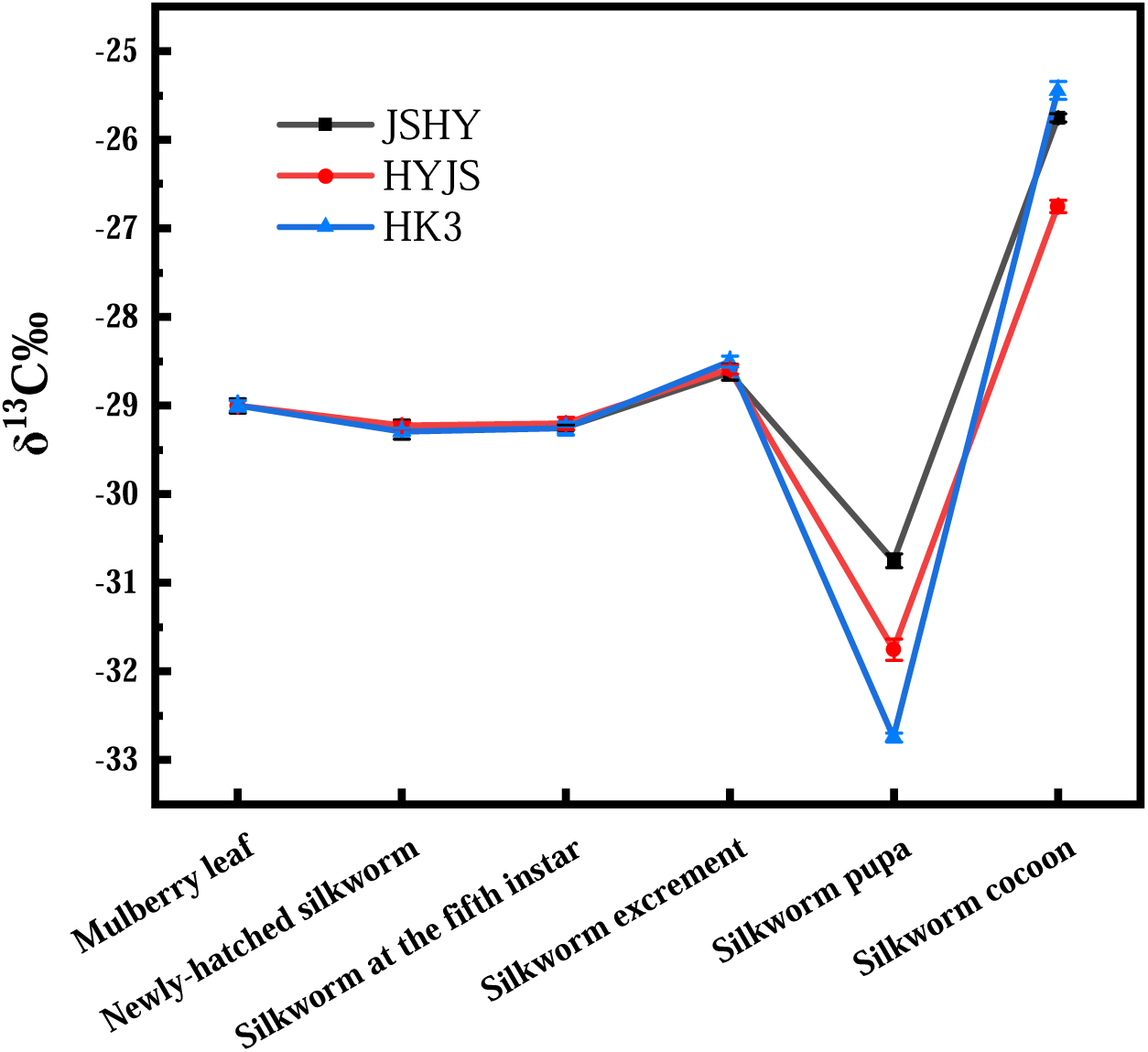
Line chart of carbon stable isotope ratios in samples

The fractionation of nitrogen, depicted in Figure 2, is more complicated than that of hydrogen. The δ^15^N values from the 3 silkworm strains differed significantly in certain samples, including in newly-hatched larvae, but not in silkworm excrement or fifth-instar larvae samples. The significant differences of δ^15^N among the 3 newly-hatched silkworm strains might be attributed to the previous generation and indicates that mating and egg laying may result in an inconsistent kinetic nitrogen isotope fractionation among the different strains, which may selective release ^14^N into sperm and eggs (Mizota and Yamanaka, 2011). The same results were found for δ^2^H and δ^18^O. However, the difference of δ^15^N value in the fifth instar silkworm was reduced to 0.1‰. The enrichment of δ^15^N in silkworms may be due to transamination during protein synthesis (Doucett et al., 1999; Jardine et al., 2004; Minagawa and Wada, 1984). What is noteworthy is that almost all the heavy elements within the silkworm excrement were enriched, except for δ^15^N. Moreover, the δ^15^N values of silkworm cocoons varied greatly between strains. These changes may be attributed to the fact that the spinning process of a silkworm requires high energy input and involves complex biochemical reactions. At the same time, it is unclear how differential routing of amino acids might affect the fractionation of stable isotopes among the 3 silkworm strains. But what is clear, is that there are myriad opportunities during cocooning for isotopic discrimination to occur. Therefore, due to its instability, the δ^15^N value was deemed not credible to be used as a sole indicator of the origin of silk fabrics.

**Figure 2.**
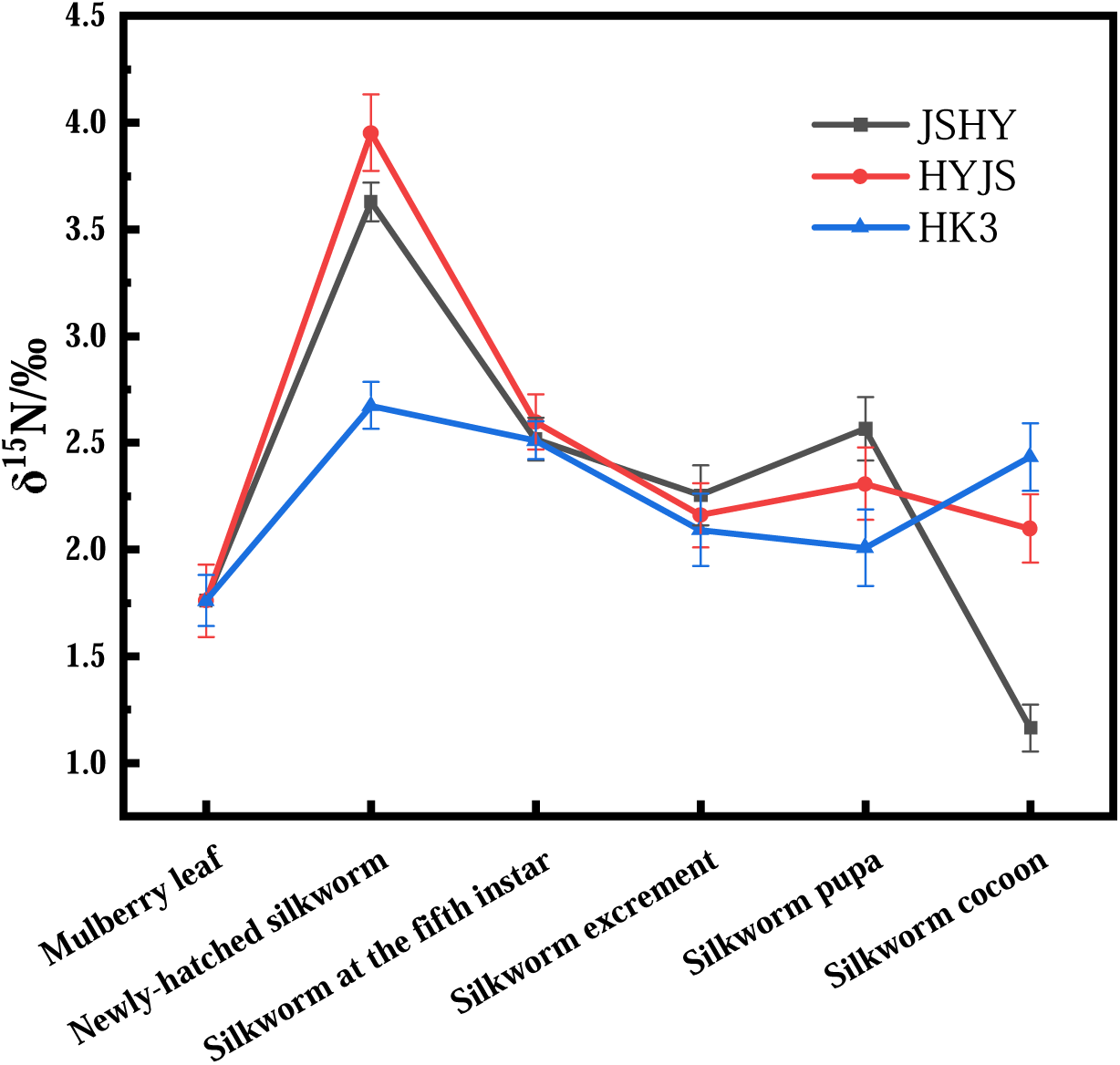
Line chart of nitrogen stable isotope ratios in samples

The δ^2^H values displayed little variation across the three different silkworm strains (Figure 3). However, the δ^2^H value of silkworm excrement samples were significantly higher than that of the larvae, which is the opposite distribution compared to that seen in δ^15^N values, suggesting an enrichment of deuterium in silkworm excrement. The δ^2^H value decreased from fifth-instar larvae to pupae but was significantly enriched in the silkworm cocoons, which was consistent with δ^13^C enrichment.

**Figure 3.**
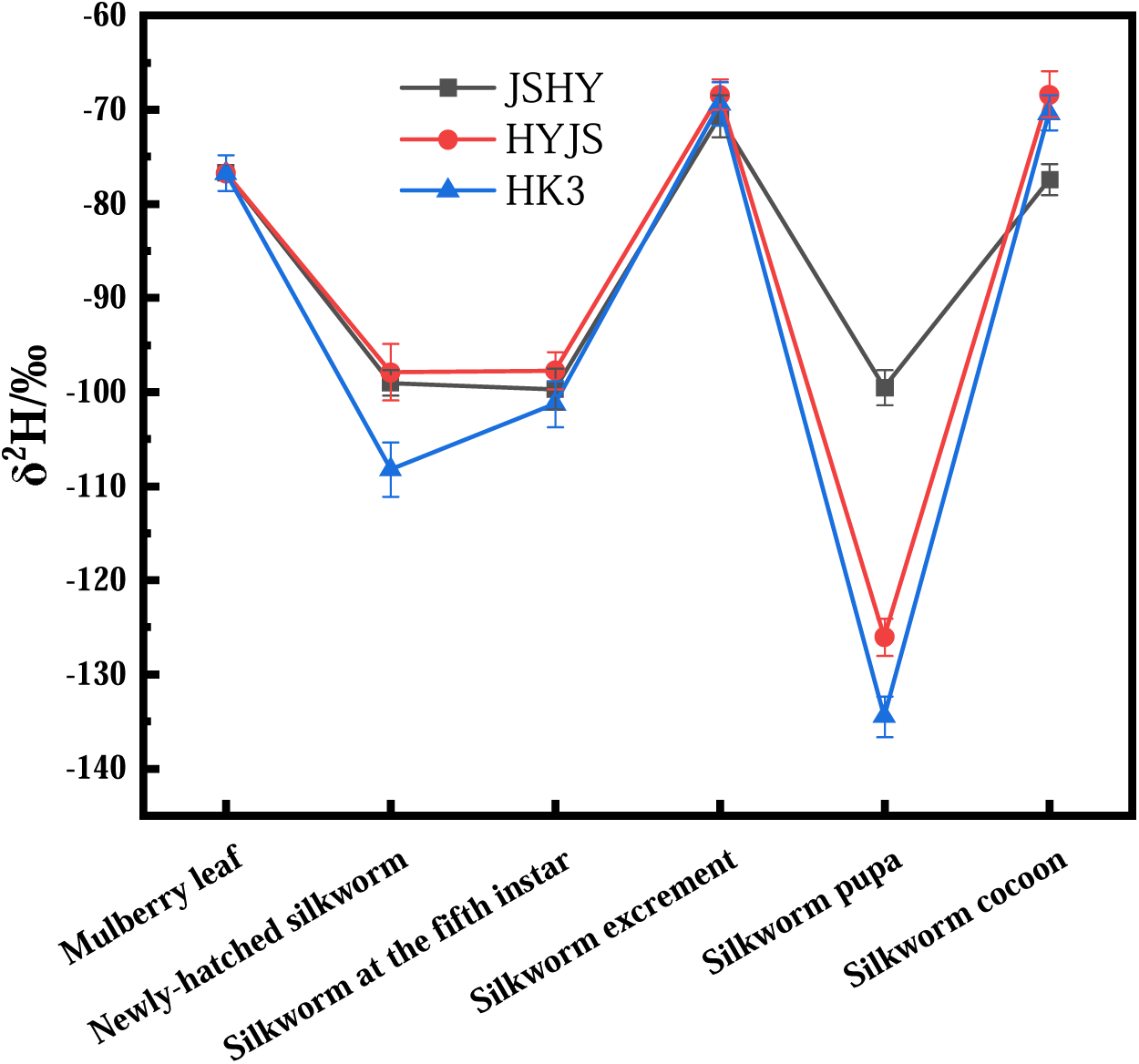
Line chart of hydrogen stable isotope ratios in samples

Finally, the δ^18^O values differed insignificantly among the three different silkworm strains and the variation across the silkworm samples was similar to that of the δ^2^H values (Figure 4). Indeed, the δ^18^O value of silkworm excrement samples was higher than that of the larvae, decreased from larva to pupae and was finally increased again in the cocoon samples, which was consistent with the variation trend of δ^2^H.

**Figure 4.**
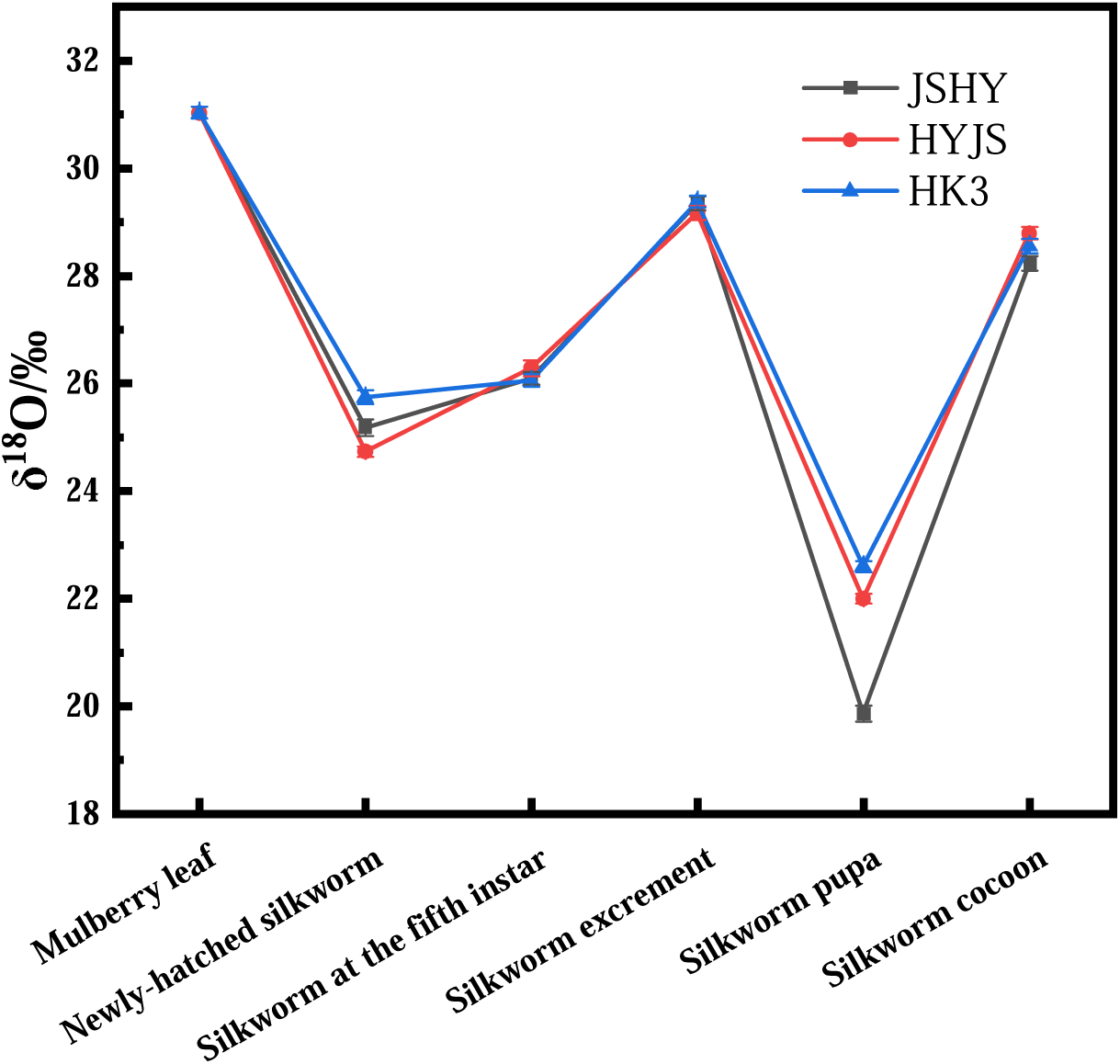
Line chart of oxygen stable isotope ratios in samples

#### Stable Isotope Fractionation of Hydrogen, Oxygen, Carbon and Nitrogen

The scatter diagram for nitrogen and carbon isotope values (Figure 5) shows that the enrichment among mulberry leaf, silkworm excrement, silkworm at fifth instar and newly-hatched silkworm samples is assumed to be low (<0.5‰). In contrast, δ^13^C was enriched in silkworm cocoons, a phenomenon likely linked to the discrimination of light and heavy isotopes derived from a combination of biochemical and physiological processes, such as metamorphosis(Doi et al., 2010; O’Brien et al., 2004; Tibbets and Rio, 2008). By contrast, the δ^15^N levels change significantly with diet. Indeed, heavy nitrogen was enriched in newly-hatched silkworms relative to the mulberry leaves they consumed. The mulberry leaf, fifth-instar larva, and newly-hatched larva samples arranged on a nearly vertical line in Figure 5, suggesting fractionation of nitrogen stable isotopes during larval development but not of carbon.

**Figure 5.**
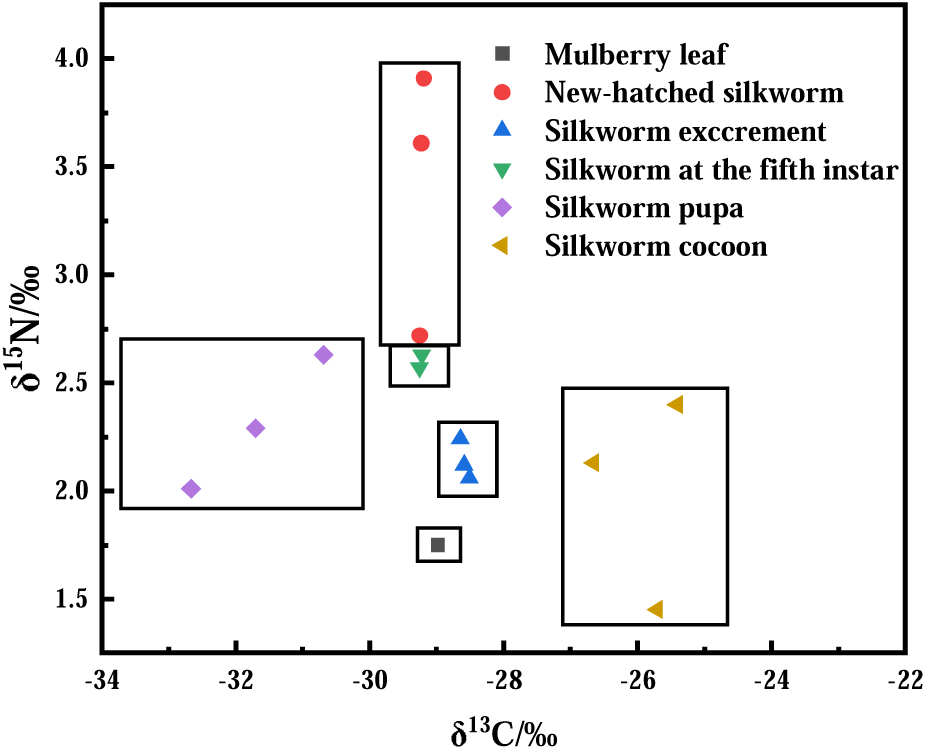
Distribution map of carbon and nitrogen stable isotope composition in samples

The scatter diagram for hydrogen and oxygen (Figure 6) revealed a distinct separation in the clustering of different silkworm samples. Silkworm pupae had the lowest δ^2^H and δ^18^O values, suggesting the depletion of hydrogen and oxygen stable isotopes in silkworm pupae. The distributions of hydrogen and oxygen stable isotopes in newly-hatched larvae and silkworm at fifth instar were found to be small, compact and concentrated, suggesting the ratios are not helpful in distinguishing instars. Mulberry leaf and silkworm excrement samples both displayed the largest δ^2^H and δ^18^O values. Overall, these data reveal that the hydrogen and oxygen stable isotopes undergo significant fractionation during the lifetime of *Bombyx mori*.

**Figure 6.**
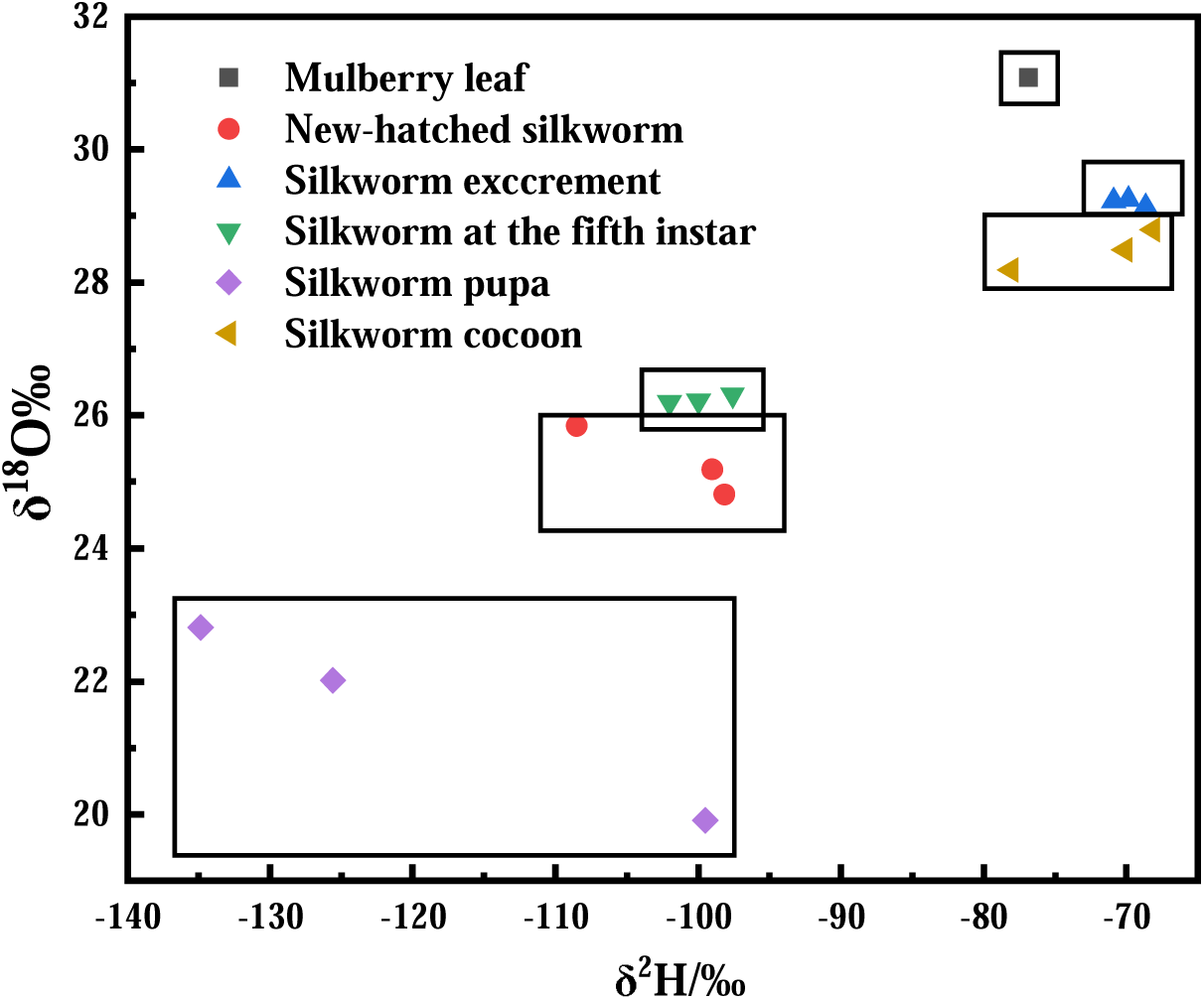
Distribution map of hydrogen and oxygen stable isotope composition in samples.

#### The effect of starvation stress on the isotope composition ratios during silkworm development

The results of stable isotope values of *Bombyx mori* under different starvation conditions are shown in Figure 7. It is obvious from the data that the δ^13^C values were significantly enriched in silkworms after starvation and there was a statistically significant positive correlation between the isotope values and starvation time. Moreover, with increasing starvation time, the δ^13^C values of both JSHY and HYJS silkworms increased by approximately 0.6‰, indicating that both stains of silkworms tend to use ^13^C under hunger stress. The different pattern of δ^18^O values for both strains with time of starvation showed an opposite trend to δ^13^C. On the other hand, the δ^15^N values of JSHY silkworms increased 0.8‰, while the δ^15^N values of HYJS silkworms decreased by 0.8‰ over 72 hours. Furthermore, the changes in heavy nitrogen levels with increasing starvation time followed an opposite trend in the two different strains, with the highest variation observed at 24 hours of starvation. This indicates that different silkworm stains of silkworm exhibit different nitrogen isotope usage under hunger stress.

**Figure 7.**
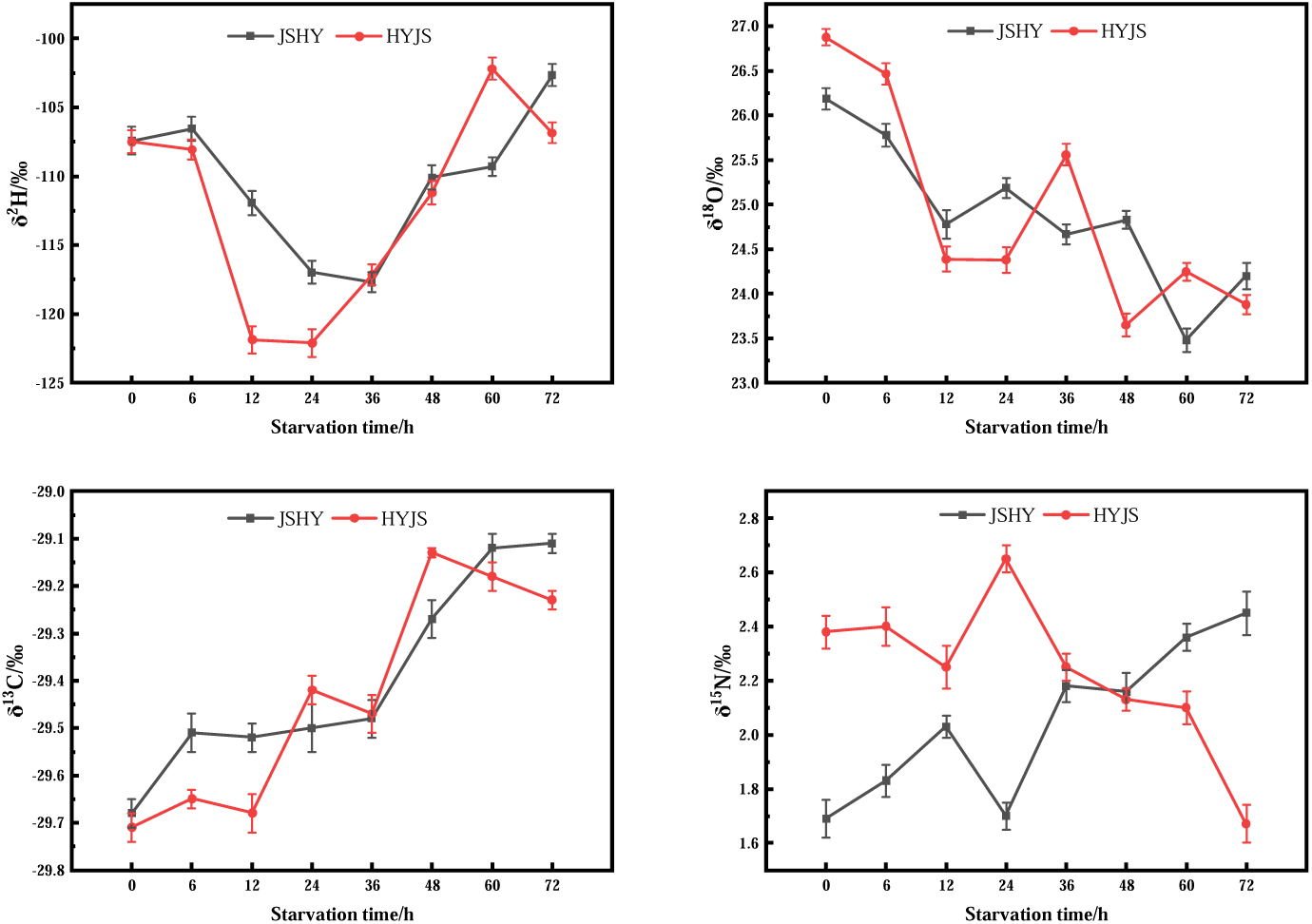
Distribution of stable hydrogen, oxygen, carbon, and nitrogen isotope compositions under starvation conditions.

The stable hydrogen, oxygen, carbon, and nitrogen isotope ratios were compared before and after three days of starvation and after three days of re-feeding (Figure 8). After resumption of a normal diet, the δ^13^C values of the silkworms continued to be enriched, indicating that the fractionation of δ^13^C is mainly influenced by foreign nutrients and metabolism rather than starvation pressure. The same results were seen for δ^2^H. However, the δ^18^O values of both silkworm strains were decreased after 72 h of starvation but returned to original levels after resumption of diet. These results suggest a statistically significant diet-dependent change in the δ^18^O values. Conversely, the analysis of nitrogen stable isotope levels across the two strains revealed significant strain-specific differences after three days of re-feeding. This suggests that there is a differential selection of ^15^N by JSHY and HYJS strains under starvation stress.

**Figure 8.**
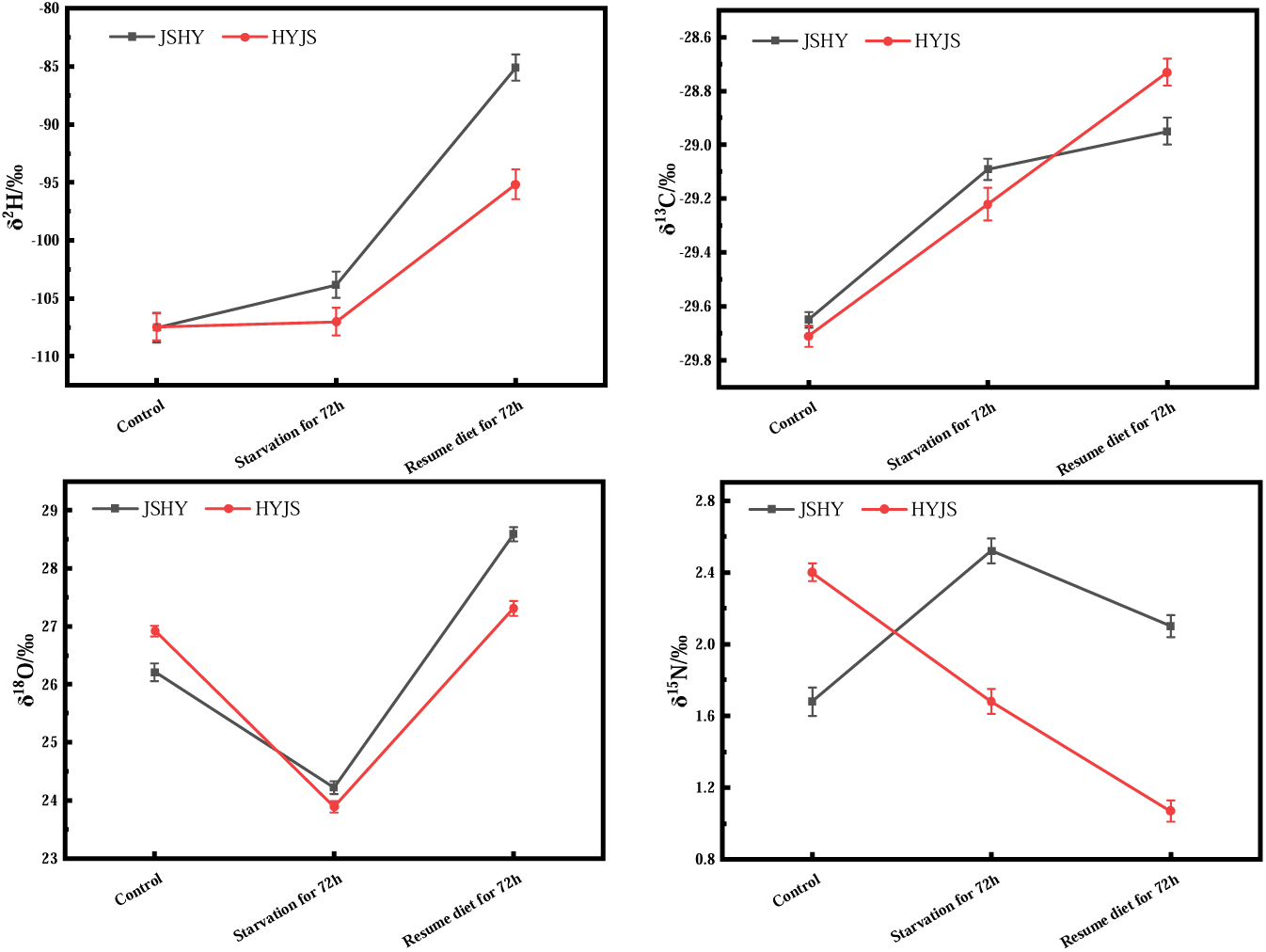
Changes of stable isotopes of hydrogen, oxygen, carbon, and nitrogen before and after starvation

#### Discriminant Analysis of Hydrogen, Oxygen, Carbon and Nitrogen Stable Isotopes

LDA, first proposed by Fisher (Pan et al., 2007), is a supervised pattern recognition method (Liu et al., 2020) carried out in an m-space (m=number of variables). It calculates an m-1 dimensional surface that separates two established categories as far as possible. According to the criteria of minimizing intra-class variance and maximizing inter-class variance, a discriminant function is obtained and then used to classify the discriminant samples (Zhang et al., 2012). The mulberry leaf, newly-hatched larva, fifth-instar larva, excrement, pupae and cocoon samples were used for discriminant analysis. The model was based on four independent variables including δ^2^H, δ^18^O, δ^13^C, and δ^15^N and the results were shown in Figure 9. The discriminant score analysis (Table 1) revealed the respective classification accuracy of the back generation and the cross-examination of δ^2^H, δ^18^O, δ^13^C and δ^15^N of samples to be 100.0% and 100.0% respectively, indicating that each of the seven sample types had a distinct stable isotope “fingerprint.”

**Figure 9.**
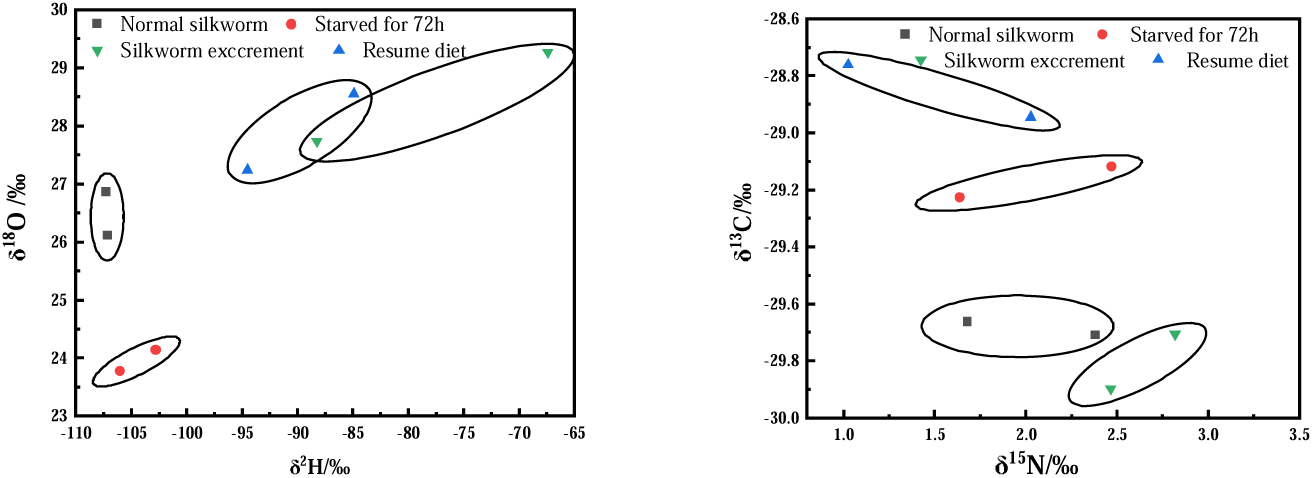
Distribution of stable isotopes of hydrogen, oxygen, carbon and nitrogen before and after starvation.

**Figure 10.**
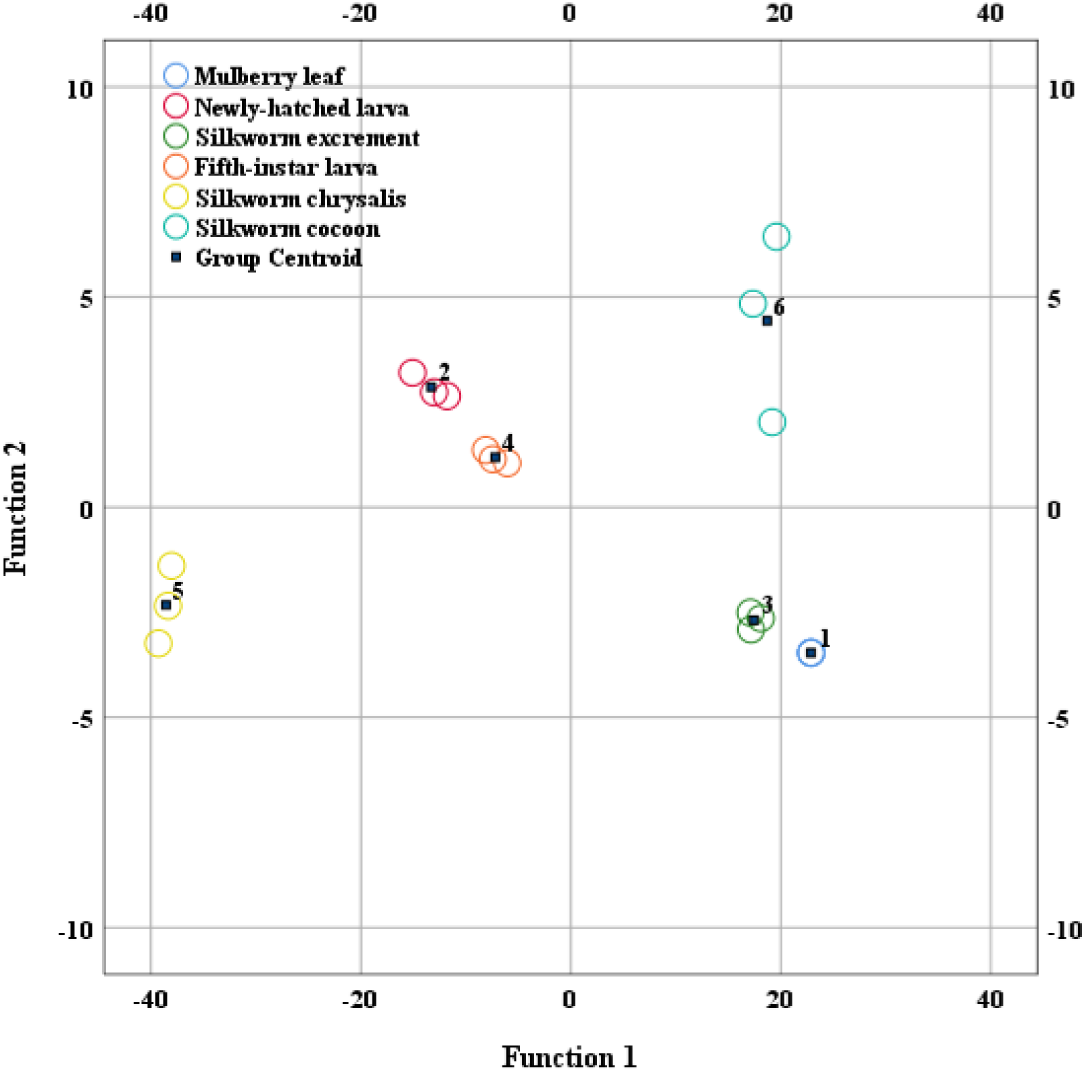
Discriminant analysis of samples.

**Table 1.** Discrimination results of samples.

**Classification Results<sup>a,c</sup>**
|  |  | Number | Predicted Group Membership |  |  |  |  |  | Total |
| --- | --- | --- | --- | --- | --- | --- | --- | --- | --- |
|  |  |  | 1 | 2 | 3 | 4 | 5 | 6 |  |
| Original | Count | 1 | 3 | 0 | 0 | 0 | 0 | 0 | 3 |
|  |  | 2 | 0 | 3 | 0 | 0 | 0 | 0 | 3 |
|  |  | 3 | 0 | 0 | 3 | 0 | 0 | 0 | 3 |
|  |  | 4 | 0 | 0 | 0 | 3 | 0 | 0 | 3 |
|  |  | 5 | 0 | 0 | 0 | 0 | 3 | 0 | 3 |
|  |  | 6 | 0 | 0 | 0 | 0 | 0 | 3 | 3 |
|  | % | 1 | 100.0 | 0 | 0 | 0 | 0 | 0 | 100.0 |
|  |  | 2 | 0 | 100.0 | 0 | 0 | 0 | 0 | 100.0 |
|  |  | 3 | 0 | 0 | 100.0 | 0 | 0 | 0 | 100.0 |
|  |  | 4 | 0 | 0 | 0 | 100.0 | 0 | 0 | 100.0 |
|  |  | 5 | 0 | 0 | 0 | 0 | 100.0 | 0 | 100.0 |
|  |  | 6 | 0 | 0 | 0 | 0 | 0 | 100.0 | 100.0 |
| Cross-validated<sup>b</sup> | Count | 1 | 3 | 0 | 0 | 0 | 0 | 0 | 3 |
|  |  | 2 | 0 | 3 | 0 | 0 | 0 | 0 | 3 |
|  |  | 3 | 0 | 0 | 3 | 0 | 0 | 0 | 3 |
|  |  | 4 | 0 | 0 | 0 | 3 | 0 | 0 | 3 |
|  |  | 5 | 0 | 0 | 0 | 0 | 3 | 0 | 3 |
|  |  | 6 | 0 | 0 | 0 | 0 | 0 | 3 | 3 |
|  | % | 1 | 100.0 | 0 | 0 | 0 | 0 | 0 | 100.0 |
|  |  | 2 | 0 | 100.0 | 0 | 0 | 0 | 0 | 100.0 |
|  |  | 3 | 0 | 0 | 100.0 | 0 | 0 | 0 | 100.0 |
|  |  | 4 | 0 | 0 | 0 | 100.0 | 0 | 0 | 100.0 |
|  |  | 5 | 0 | 0 | 0 | 0 | 100.0 | 0 | 100.0 |
|  |  | 6 | 0 | 0 | 0 | 0 | 0 | 100.0 | 100.0 |
| a. 100.0% of original grouped cases are correctly classified. |  |  |  |  |  |  |  |  |  |
| b. Cross validation is done only for those cases in the analysis. In cross validation, each case is classified by the functions derived from all cases other than that case. |  |  |  |  |  |  |  |  |  |
| c. 100.0% of cross-validated grouped cases are correctly classified. |  |  |  |  |  |  |  |  |  |
| 1 – Mulberry leaf. 2 – Newly-hatched larva. 3 - Excrement. 4 – Fifth-instar larva. 5. Pupa. 6 – Cocoon. |  |  |  |  |  |  |  |  |  |

## Conclusion

Silkworm strain had little effect on the carbon stable isotope ratios in the majority of the silkworm-related samples. A large difference in δ^15^N was observed within newly-hatched silkworm samples in JSHY and HK3 strains, but the ^15^N levels in the fifth instar silkworms and their excrement was relatively stable. The difference of δ^15^N among the three newly-hatched silkworm strains indicated that mating and egg laying may result in an inconsistent kinetic nitrogen isotope fractionation. In contrast to δ^15^N, the δ^2^H and δ^13^C values were enriched in silkworm cocoon samples. Moreover, the δ^13^C signatures of silkworm samples displayed significant starvation-time effects, whereas the δ^15^N values displayed a diametrically opposite trend between the JSHY and HYJS strains. This indicates that different silkworm strains exhibit different nitrogen isotope usage under starvation stress. The carbon stable isotope ratios in silkworm excrement, fifth-instar larvae, and pupae samples demonstrated significant fractionation, while the newly-hatched larvae, fifth instar larvae, and mulberry leaf samples exhibited mostly nitrogen fractionation. Mulberry leaf and silkworm excrement were found to be highly enriched for ^2^H and ^18^O.

Starvation experiments revealed that δ^13^C levels increased due to metabolic requirements as indicated by continual enrichment of δ^13^C after diet resumption for 72 h. Similarly, heavy hydrogen was found to be gradually enriched under starvation and increased further upon diet resumption. However, ^18^O was depleted after starving and enriched after re-feeding, which indicated that starvation greatly impacts δ^18^O, but not δ^2^H. Under starvation conditions, orthogonal (JSHY) and anti-crossing (HYJS) newly-hatched silkworms accumulated heavy nitrogen differently, showing that the different hybridization methods had a great impact on the larval feeding habits. Meanwhile, isotopic fractionation occurred in the silkworm ecological process.

Finally, LDA was able to discriminate the mulberry leaf, newly-hatched larvae, silkworm excrement, fifth-instar larvae, silkworm pupae, and silkworm cocoons with a classification accuracy rate of 100.0%, indicating that the specific “fingerprint” information carried by stable isotopes can effectively be utilized to identify and distinguish each of these life stages and substances. Due to the high classification accuracy rate and the appropriate discrimination elements, we propose that IRMS can be used to help trace nutrient transmission during silkworm development and silk production. As a result, the combination of LDA and isotope measurements may provide the foundation for new techniques and strategies for the traceability of textiles.

## Acknowledgments

We thank Sericulture Institute of Zhejiang Academy of Agricultural Sciences for providing the experimental materials.

## Authors’ contributions

Conception and design of study: Zhiqin Peng, Yang Zhou.

Acquisition of data: Hao Li, Yujie He, Jingzhong Lu, Yong Liu, Dan Yang, Shuai Shao, Gang Lv, Hailiang Yang, Hailing Zheng, Yang Zhou.

Analysis and interpretation of data: Hao Li, Yujie He, Jingzhong Lu, Zhiqin Peng, Liling Jia, Xuhong Cui.

Drafting the manuscript: Hao Li, Yujie He, Zhiqin Peng.

Revising the manuscript critically for important intellectual content: Hao Li and Zhiqin Peng.

The authors agree to take responsibility for all aspects of the work to ensure that questions relating to the accuracy or completeness of any part of the work are properly investigated and resolved.

